# Bioinf-Farma: supervised integration of epitope prediction and recombinant protein developability for automated vaccine candidate prioritization

**DOI:** 10.64898/2026.06.15.732271

**Authors:** Heather Bondi, Matteo Crespi, Marco Orlando, Francesco Lescai, Stefano A. Serapian, Giorgio Colombo, Mauro Fasano, Loredano Pollegioni, Gianluca Molla

**Affiliations:** Department of Science and High Technology, University of Insubria, Busto Arsizio, Italy; Department of Biotechnology and Life Sciences, University of Insubria, Varese, Italy; Laboratory for Environmental and Life Sciences, University of Nova Gorica, Slovenia; Department of Biology and Biotechnology, University of Pavia, Pavia, Italy; Department of Chemistry, University of Pavia, Pavia, Italy

## Abstract

Vaccine antigen discovery requires prioritizing protein candidates according to both immunogenic potential and recombinant expression feasibility. These properties are typically evaluated using separate computational tools, requiring researchers to integrate heterogeneous outputs through ad hoc workflows. Here, we present BIOINF-farma, a modular platform integrating epitope prediction and developability assessment for rational antigen selection within a unified environment. Candidates can be submitted as amino acid sequences or three-dimensional structures. When experimental structures are unavailable, BIOINF-farma automatically searches for models in AlphaFold DB or performs structure prediction using Boltz-2, ensuring a standardized structural representation for downstream analyses. Antigenicity is quantified by combining structure-based conformational epitope signals (MLCE/REBELOT-BEPPE) and sequence-based linear epitope propensity scores (BepiPred 3.0) into a protein-level Antigenicity Score, with a classification threshold optimized on a manually curated validation dataset. Developability is evaluated through two supervised Random Forest meta-learners that integrate three solubility predictors (DeepSoluE, SoluProt, Protein–Sol) and three thermal stability predictors (TemStaPro, ProLaTherm, BertThermo), whose outputs are combined into an Expression Efficiency Score (EES). By integrating complementary predictive signals, the meta-learning framework achieves greater accuracy and robustness than individual predictors while maintaining performance across a broad range of sequence identities. The Antigenicity Score effectively discriminates antigenic from non-antigenic proteins with a large effect size, whereas EES successfully distinguishes soluble from insoluble outcomes on an independent panel of recombinant proteins expressed in *Escherichia coli*. BIOINF-farma jointly assesses antigenicity and expression feasibility within a single framework. Its modular architecture facilitates the incorporation of future predictive methods, while its web-based interface makes the full pipeline accessible to users without programming expertise, supporting rapid candidate triage in vaccine research and emerging pathogen responses.

**Author Summary:** Vaccine development begins with a critical step: identifying, among the many proteins encoded in a pathogen genome, those most suitable as candidate antigens. A promising candidate must satisfy two requirements that are rarely evaluated together. It must be recognized by the immune system, so that vaccination elicits a protective response; and it must be amenable to recombinant production, since antigens that cannot be obtained in sufficient quantity and quality are of limited practical use. Current computational tools typically address only one of these aspects, and researchers must integrate their outputs manually, through procedures that are time-consuming and prone to inconsistency. We developed BIOINF-farma, an automated platform that brings these two assessments into a single analytical framework. Starting from a protein sequence or an experimental structure, the platform retrieves or predicts a three-dimensional model, evaluates the protein’s antigenic potential by combining complementary epitope predictors, and estimates its expression feasibility by integrating multiple solubility and stability predictors through supervised machine learning. A web-based interface makes the full workflow available to experimental immunologists and vaccine developers without requiring computational expertise, supporting rational candidate prioritization in routine vaccine research and during emerging pathogen responses.

## Introduction

The identification of suitable antigens is a necessary step prior to the development of recombinant vaccines and immunotherapeutic strategies against emerging pathogens. This process relies on the rapid generation and sharing of genomic and proteomic data from newly identified strains. Recent viral outbreaks, such as Zika virus and SARS-CoV-2, have demonstrated the ability of the scientific and clinical communities to promptly generate and disseminate large-scale omics datasets [1,2]. Advances in high-throughput sequencing, structural biology, and computational methods have enabled the rapid production and analysis of these datasets.

However, the availability of large-scale datasets alone is insufficient. Rather, it shifts the bottleneck downstream, where efficient and reliable predictive systems are required to identify antigenic proteins and prioritize candidates suitable for vaccine development. In this context, the adoption of advanced computational approaches, including machine learning (ML) algorithms, can accelerate antigen discovery and support the development of novel vaccines. These approaches are inherently faster and less resource-intensive than traditional experimental strategies [3–5]. Computational predictors are widely used to estimate epitope content, antigenicity, and relevant physicochemical properties of viral and bacterial proteins, thereby reducing the number of experimental validation steps, which are often time- and resource-consuming. Nevertheless, despite significant methodological progress, the computational identification of effective antigens remains challenging. A major limitation of current computational approaches is that antigenicity prediction is typically uncoupled from the assessment of recombinant protein producibility, including solubility and stability in heterologous systems such as *Escherichia coli*. Consequently, candidates selected solely on predicted immunogenicity often display poor recombinant yield and require extensive experimental optimization [6,7].

A further limitation of current computational approaches is the relatively low reliability of individual prediction tools when applied to antigen discovery. This limitation does not depend solely on model accuracy, but also reflects the heterogeneous strategies used to identify antigenic determinants.

Sequence-based predictors typically estimate linear B-cell epitope propensity and antigenicity scores using local amino acid information [8,9]. In contrast, conformational epitopes are identified through structure-based approaches that evaluate surface accessibility, residue energetics, and spatial clustering [10,11].

Because these methods target different types of antigenic determinants, rely on distinct input data, and employ different algorithms, their predictions are often complementary but only partially overlapping. This heterogeneity often results in inconsistent antigen mapping when predictors are used independently.

In parallel, the prediction of physicochemical properties underlying efficient recombinant protein expression remains challenging. Solubility, aggregation propensity, and stability depend on protein folding, three-dimensional structure, and interactions with the heterologous host cellular environment [12]. Nevertheless, many predictors infer these properties from primary sequence alone, often resulting in reduced generalizability outside the distributions represented in their training datasets.

Collectively, these limitations highlight the need for integrated computational frameworks capable of jointly evaluating antigenicity and recombinant protein production during vaccine antigen prioritization. Recent advances in ML have enabled the integration of heterogeneous descriptors derived from sequence-, structure-, and physicochemical-based predictors, combining antigenicity estimators with solubility and stability predictors within unified frameworks. By learning the relative contribution of different features directly from experimental data, this strategy reduces dependence on predefined weights or user-defined heuristics and improves the robustness of candidate prioritization.

Here, we present Bioinf-Farma, an automated, modular, and extensible computational platform that integrates protein structure retrieval or prediction from submitted sequences, linear and conformational epitope prediction, and solubility and stability assessment. Central to this framework is the use of supervised machine learning models trained on curated datasets to integrate heterogeneous antigenicity and producibility descriptors into unified predictive scores, thereby improving the robustness of candidate prioritization. Collectively, these components establish Bioinf-Farma as an accessible and flexible platform for rational antigen selection in vaccine and immunotherapy research.

### Design and implementation

#### Overview of the BIOINF-farma platform

BIOINF-farma is a modular computational platform designed to jointly evaluate antigenicity and recombinant protein producibility during antigen candidate prioritization. Its design addresses a major limitation in current antigen discovery workflows, in which epitope prediction and recombinant expression assessment are typically performed using separate and poorly integrated tools. The platform is organized into four sequential modules (Fig 1): (i) a protein 3D structure preparation module, which retrieves experimental structures or predicts three-dimensional models for each input sequence; (ii) an antigenicity module, which integrates structure-based and sequence-based epitope predictors into a single protein-level Antigenicity Score; (iii) a developability module, which combines solubility and thermal stability predictions through supervised meta-learning into an Expression Efficiency Score (EES); and (iv) a reporting module, which organizes individual and integrated predictor scores into structured outputs. Each candidate protein is processed independently, enabling execution of large-scale input datasets. Inputs are accepted either as amino acid sequences (FASTA) or as three-dimensional structures (PDB); intermediate outputs are standardized across modules to ensure interoperability, and final outputs are aggregated at both residue and protein level. A browser-based graphical interface (see Web-based user interface) enables remote access to the full pipeline without requiring command-line experience or local installation.

**Fig 1.**
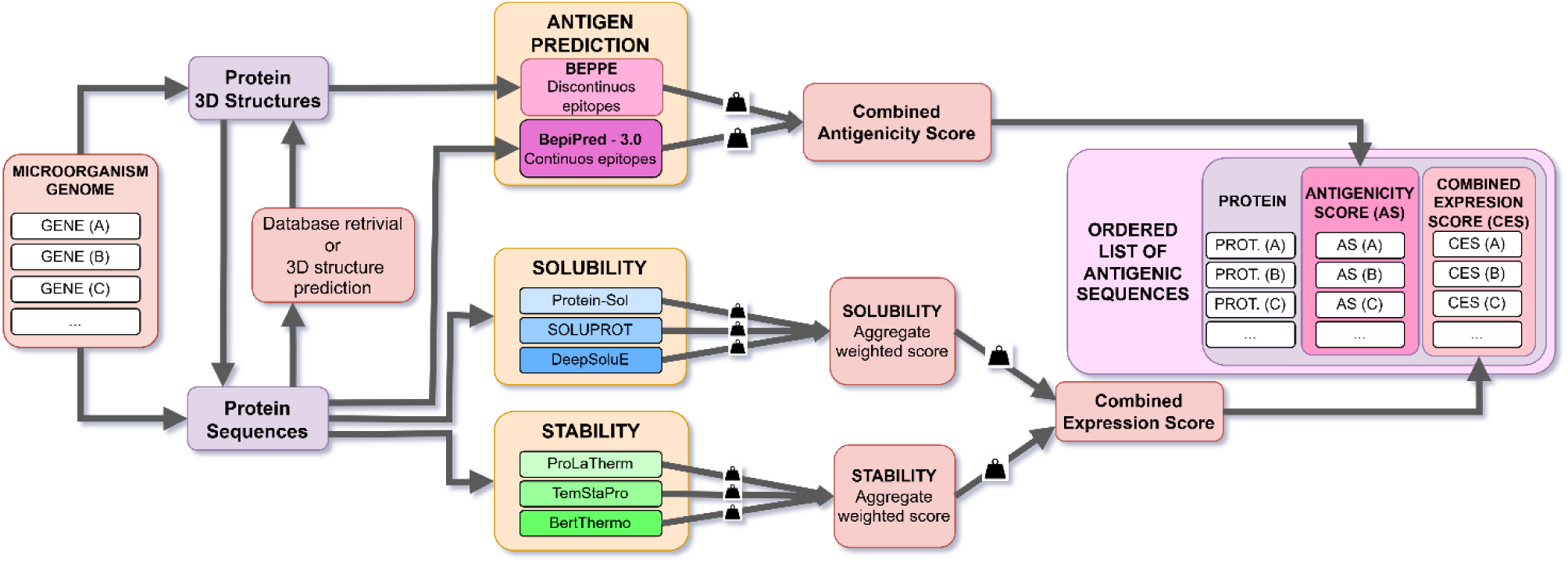
Overview of the BIOINF-farma platform architecture. Protein sequences derived from microbial genomes are processed through modular analysis components. Structural information is retrieved or predicted prior to epitope mapping, where structure-based (BEPPE) and sequence-based (BepiPred 3.0) predictions are integrated into a combined Antigenicity Score. In parallel, solubility (Protein-Sol, SoluProt, DeepSoluE) and thermal stability (ProLaTherm, TemStaPro, BertThermo) predictors are aggregated into weighted developability scores. Solubility and stability are further integrated into a combined Expression Efficiency Score, enabling ranking of candidate antigenic proteins based on both immunogenic potential and developability properties.

#### Structure retrieval and prediction

When protein candidates are submitted as amino acid sequences, BIOINF-farma retrieves or produces the three-dimensional structural models required for conformational epitope mapping. The system prioritizes retrieval of experimental structures when available, and, when homologous structures are unavailable, generates accurate computational structural models.

#### Homology-based structure retrieval

For each submitted sequence, a (either remote or local) database of experimental (PDB) or predicted (AlphaFold-DB) protein structures is first searched using hhsearch [13] and mmseqs2 [14], respectively. If a 99% identity and coverage match is found, the matched experimental (https://www.rcsb.org/) or AlphaFold-DB (https://alphafold.ebi.ac.uk/) structure is retained.

#### De novo structure prediction

When homologous structures cannot be identified, a three-dimensional structure is predicted from sequence using Boltz-2 [15] in MSA and template mode. Five diffusion samples are generated with the “--use_potentials” inference option enabled. The model with the highest confidence_score is selected for downstream evaluation. Predicted structures are retained only if the selected model exhibits an average pLDDT > 0.75 and a confidence_score > 0.8; accepted models are subsequently converted to PDB format for downstream analyses.

#### Structure standardization and sequence extraction

Protein structures are preprocessed with pdb4amber (AmberTools23) [16] to ensure consistency with downstream structure-based analyses: this includes removal of heteroatoms, resolution of alternate conformations, and addition of missing heavy atoms, yielding standardized and Amber-compatible structures. Amino acid sequences are reconstructed from Cα atom records and converted from three-letter to one-letter residue codes. The resulting sequences are exported in FASTA format and used as input for all sequence-based prediction tools.

#### Antigenicity module: integrating structure- and sequence-based epitope signals

The antigenicity module was designed to quantify the antigenic potential of candidate proteins by integrating complementary structure- and sequence-based B-cell epitope predictors within a unified framework.

#### Structure-based epitope prediction

Structure-based prediction of conformational epitopes is performed using the energy decomposition strategy implemented in the MLCE approach, which identifies epitopes by detecting weakly coupled surface residue patches on the three-dimensional protein structures [17,18]. This method enables the identification of conformational epitopes emerging from the native protein fold [18]. Predictions are binarized such that BEPPE(i)=1 for residues belonging to predicted patches and 0 otherwise.

#### Sequence-based epitope prediction

Sequence-based prediction of linear epitopes is performed using BepiPred 3.0, which assigns residue-level epitope scores based on sequence-derived features [19]. Predictions are binarized such that BepiPred(i)=1 for residues exceeding the 0.15 threshold and 0 otherwise.

#### Integration and Antigenicity Score calculation

Outputs from MLCE/REBELOT-BEPPE and BepiPred 3.0 are integrated at the residue level to compute a combined antigenicity score. Binary encoding was adopted to harmonize heterogeneous predictor outputs and reduce dependence on predictor-specific score distributions.

For each residue *i*, the antigenicity score is defined as:

[uamth1]

The protein-level Antigenicity Score is then computed as the arithmetic mean across all residues:

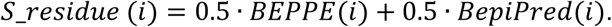

where *N* denotes the total number of residues in the protein sequence.

The resulting score is normalized from 0 to 1, where 0 indicates absence of predicted antigenic regions and 1 indicates maximal predicted antigenicity. Proteins with *S_protein* ≥ 0.417 are classified as antigenic, whereas proteins with lower scores are classified as non-antigenic. The classification threshold was determined through ROC-based optimization on the reference dataset.

#### Developability module: meta-learning integration of solubility and stability predictors

The developability module is designed to estimate recombinant protein producibility by integrating complementary sequence-based predictors of solubility and thermal stability. Rather than relying on individual predictor outputs, supervised meta-learning was used to improve robustness and reduce dependence on predictor-specific biases.

#### Solubility meta-learner

Solubility scores generated by DeepSoluE [20], SoluProt [21], and Protein–Sol [22] are integrated using a supervised Random Forest (RF) meta learner trained on experimentally annotated solubility datasets. The resulting model produces a unified solubility score, *S_solubility*, ranging from 0 to 1. Proteins with *S_solubility* ≥ 0.5 are classified as soluble, whereas proteins with lower scores are classified as insoluble.

#### Stability meta-learner

Thermal stability predictions generated by TemStaPro [23], ProLaTherm [24], and BertThermo [25] are integrated using an RF meta-learner trained on experimentally annotated thermostability datasets. The resulting model produces a unified stability score *S_stability*, ranging from 0 to 1. Proteins with *S_stability* ≥ 0.5 are classified as thermostable, whereas proteins with lower scores are classified as thermolabile.

#### Integration of solubility and stability module

Solubility and stability predictions are integrated into a single developability metric termed Expression Efficiency Score (EES). For each protein, the EES is calculated as:

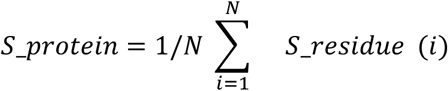

Weights were heuristically assigned to prioritize solubility, which is generally considered the principal limiting factor in recombinant protein production in *E. coli*, while retaining thermal stability as a complementary developability descriptor.

#### Software architecture

BIOINF-farma was designed as a modular computational framework for automated epitope prediction and protein developability assessment. The platform was developed and tested on Ubuntu 20.04.6. Analysis modules were implemented in Python (v3.12), whereas workflow orchestration and execution management were handled through Bash scripting.

Each module is executed sequentially with standardized input and output files to ensure interoperability across workflow stages. To guarantee reproducibility and avoid dependency conflicts among third-party bioinformatics tools, each external predictor is executed within a dedicated Conda environment. This strategy ensures stable and reproducible execution of all integrated software components. Intermediate and final outputs are stored within a standardized directory structure, enabling full traceability of all processing steps and facilitating downstream inspection. Detailed installation and usage instructions are provided in S1 Text.

#### Web-based user interface

BIOINF-farma is accessible via a web interface implemented using Apache2 and PHP and deployed on the workstation hosting the pipeline. The interface accepts PDB and FASTA files and supports file upload, input validation, and pipeline execution. Following job submission, a unique job ID is generated and the workflow is executed in the background through a dedicated wrapper script. Job progress is tracked asynchronously and displayed in real time through a dynamic progress bar. Interface updates are performed every two seconds through AJAX polling of a JSON status file.

Upon completion, the interface displays a summary of the results and provides a downloadable PDF report. Raw outputs generated by individual predictors are also accessible through the interface. A detailed tutorial for web-based usage is provided in S1 Text.

## Results

### Structure- and sequence-based epitope prediction

To quantify protein antigenicity through a unified protein-level metric, we assembled a curated Antigenicity Dataset (CAD) comprising manually annotated likely antigenic (n = 115) and likely non-antigenic proteins (n = 75) spanning human, viral, and bacterial origins. Likely non-antigenic proteins consisted predominantly of human intracellular and plasma proteins, whereas likely antigenic proteins mainly included pathogen-derived antigens such as viral structural and non-structural proteins and bacterial toxins (S2 Table).

Epitope prediction was performed using complementary structure- and sequence-based approaches. Structure-based prediction was carried out using the MLCE approach implemented in REBELOT–BEPPE, which identified conformational epitope regions as discrete surface patches [17,26,27]. Sequence-based prediction was performed using BepiPred 3.0 on FASTA sequences either provided directly or extracted from protein structures, generating residue-level epitope propensity scores [19].

Residue-level predictions from the two methods were integrated into a unified antigenicity signal and aggregated into a protein-level Antigenicity Score. The resulting score distributions differed markedly between curated antigenic and non-antigenic proteins (Fig 2A).

**Fig 2.**
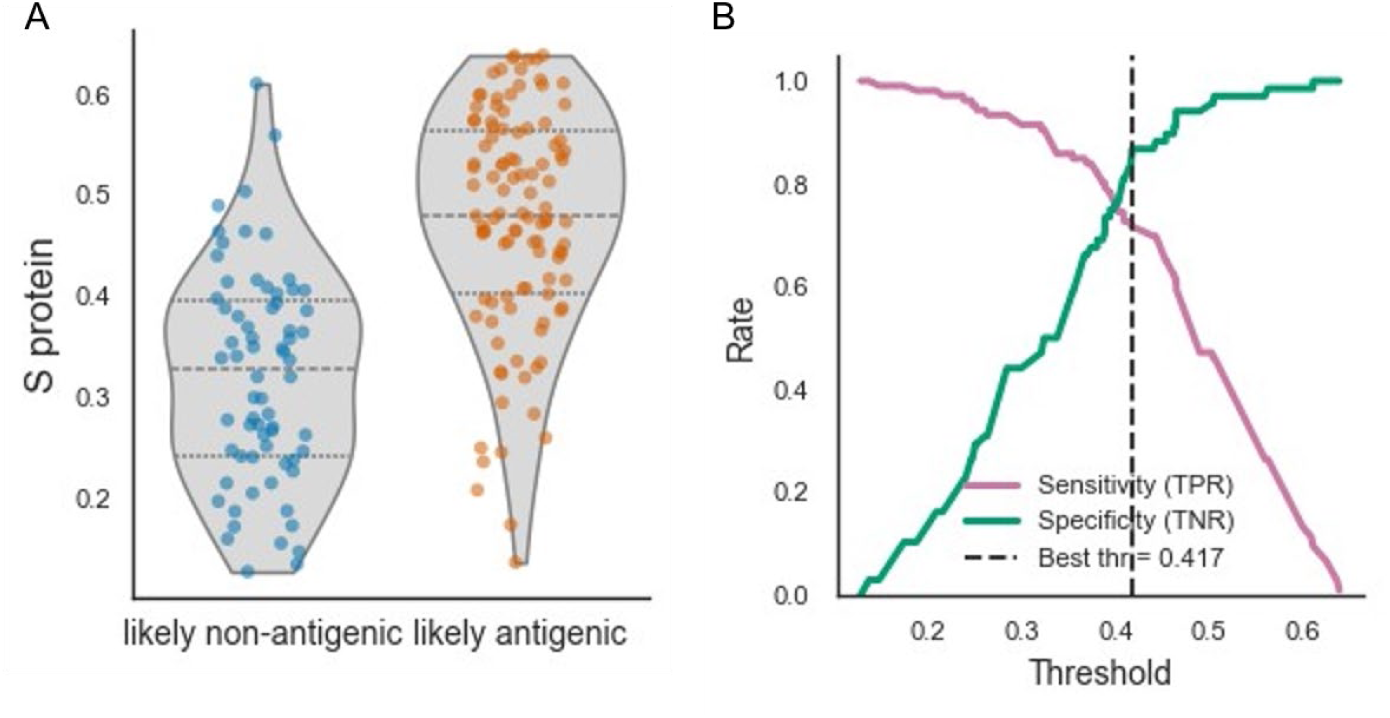
**Antigenicity score distribution and threshold selection**. (A) Distribution of the protein-level antigenicity score (S protein) across curated likely non-antigenic and likely antigenic proteins in the CAD dataset. Violin plots show score density with individual protein values overlaid. Horizontal dashed lines indicate quartiles. (B) Sensitivity (true positive rate, TPR) and specificity (true negative rate, TNR) as a function of the decision threshold. The dashed vertical line indicates the threshold selected using the Youden’s J criterion (0.417), corresponding to the balanced operating point between sensitivity and specificity.

Likely antigenic proteins showed significantly higher scores than likely non-antigenic proteins (Mann–Whitney U = 1150, p = 3.8×10⁻¹⁴), with consistent distributional differences (KS = 0.585, p = 1.068×10⁻¹³) and a large effect size (Cliff’s δ = 0.681). These results indicate that the integrated score effectively discriminates between curated antigenic and non-antigenic proteins.

A binary classification threshold was subsequently defined using the pre-specified Youden’s J criterion to balance sensitivity and specificity. This procedure identified an optimal threshold of 0.417 (Fig 2B), whereby proteins with *S_protein* ≥ 0.417were classified as antigenic and proteins with lower scores as non-antigenic.

### Developability prediction

#### Integrating solubility predictions using supervised machine learning

To build the solubility component of the EES, we first evaluated whether supervised integration of multiple predictors could improve solubility prediction performance. A large reference dataset was assembled from publicly available resources after removing all sequences used to train the original solubility prediction tools, thereby minimizing potential data leakage. The resulting dataset comprised 99,992 protein sequences, including 47,844 experimentally annotated soluble proteins and 52,148 insoluble proteins.

Sequence redundancy was assessed through all-versus-all sequence comparisons. The identity distribution revealed a heterogeneous dataset, containing both highly redundant clusters (35.49% of sequences with >90% identity) and broadly diverse sequences (Fig 3A). To reduce redundancy-related bias while preserving biological variability, redundancy reduction was performed separately for soluble and insoluble proteins. This resulted in 14,770 soluble and 19,706 insoluble non-redundant sequences (S3 Table). Sequence length distributions were subsequently examined to evaluate potential size-related biases between classes (Fig 3B).

**Fig 3.**
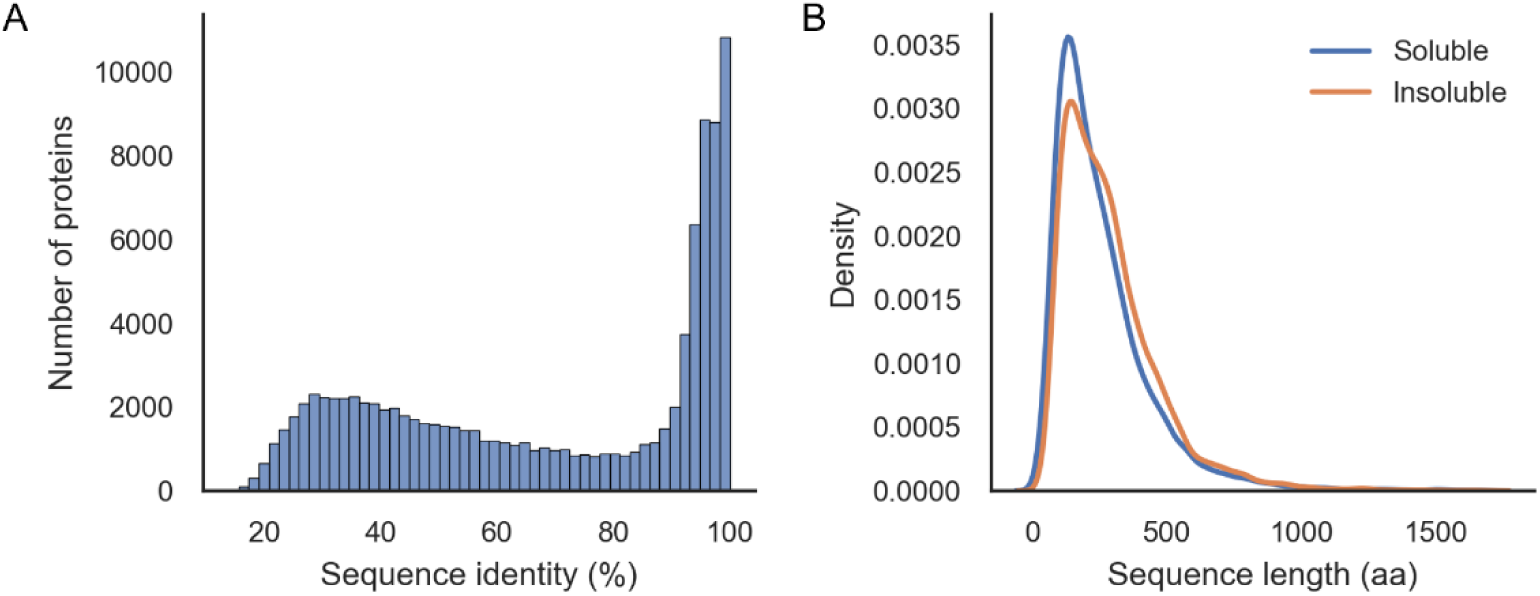
Sequence redundancy and length distribution in the curated solubility dataset. (A) Distribution of pairwise sequence identity obtained from all-versus-all comparisons, showing the proportion of sequences across identity bins. The dataset exhibits a heterogeneous structure, with a substantial fraction of highly identity sequences alongside a broad range of lower identity values. (B) Distribution of protein sequence lengths for soluble and insoluble classes. The two classes display comparable length distributions, indicating no major bias in protein size between soluble and insoluble proteins.

Solubility scores generated by DeepSoluE [20], SoluProt [21], and Protein–Sol [22] were integrated using four supervised meta-learning classifiers: Random Forest (RF), Gradient Boosting (GBC), XGBoost (XGB), and K-Nearest Neighbors (KNN). Performance was evaluated on a held-out test set, using precision as the primary optimization metric to minimize false positive predictions during antigen prioritization.

The four classifiers showed comparable performance across evaluation metrics (Table 1, Fig 4A-C). RF achieved the highest precision (0.826, 95% CI [0.805–0.846]), in line with our objective of limiting false positives. To evaluate the generalizability of this ranking beyond the optimization metric, we examined threshold-independent and threshold-dependent performance measures. RF, XGB, and GBC achieved equivalent ROC-AUC values (0.711), indicating comparable discriminatory performance across all decision thresholds, while KNN showed a marginally lower value (0.705). For Average Precision (AP), RF, XGB, and GBC achieved identical values (0.696), demonstrating equivalent ranking performance at varying thresholds, while KNN achieved a lower value (0.682). All metrics were evaluated with 95% bootstrap confidence intervals (5,000 replicates; see Table 1 for complete estimates). Based on these results, RF was selected as the solubility meta-predictor.

**Fig 4.**
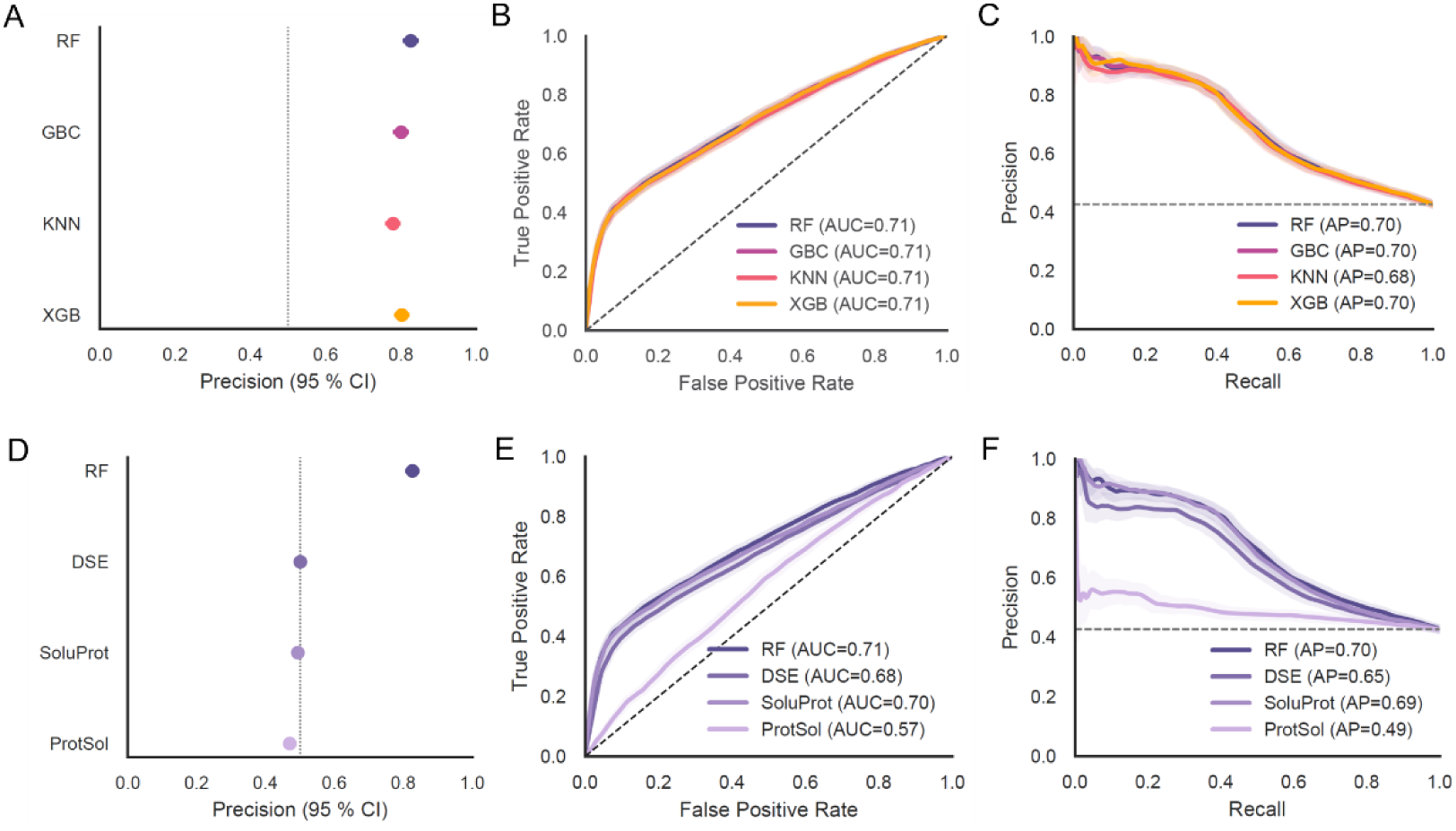
Performance of supervised meta-learning models and comparison with individual solubility predictors. **(A)** Precision values with 95% bootstrap confidence intervals for Random Forest (RF), Gradient Boosting Classifier (GBC), XGBoost (XGB), and K-Nearest Neighbors (KNN). **(B)** Receiver Operating Characteristic (ROC) curves showing threshold-independent discriminatory performance, with Area Under the Curve (AUC) values indicated. **(C)** Precision-Recall curves showing threshold-dependent ranking performance, with Average Precision (AP) values indicated.**(D)** Precision values with 95% bootstrap confidence intervals for RF and three standalone solubility predictors: DeepSoluE (DSE), SoluProt, and Protein–Sol. **(E)** ROC curves with AUC values for each method. **(F)** Precision-Recall curves with AP values for each model. Dashed lines indicate reference benchmarks: vertical (A, D: full test set precision), diagonal (B, E: random classifier, AUC=0.5), and horizontal (C, F: baseline precision).

**Table 1.**
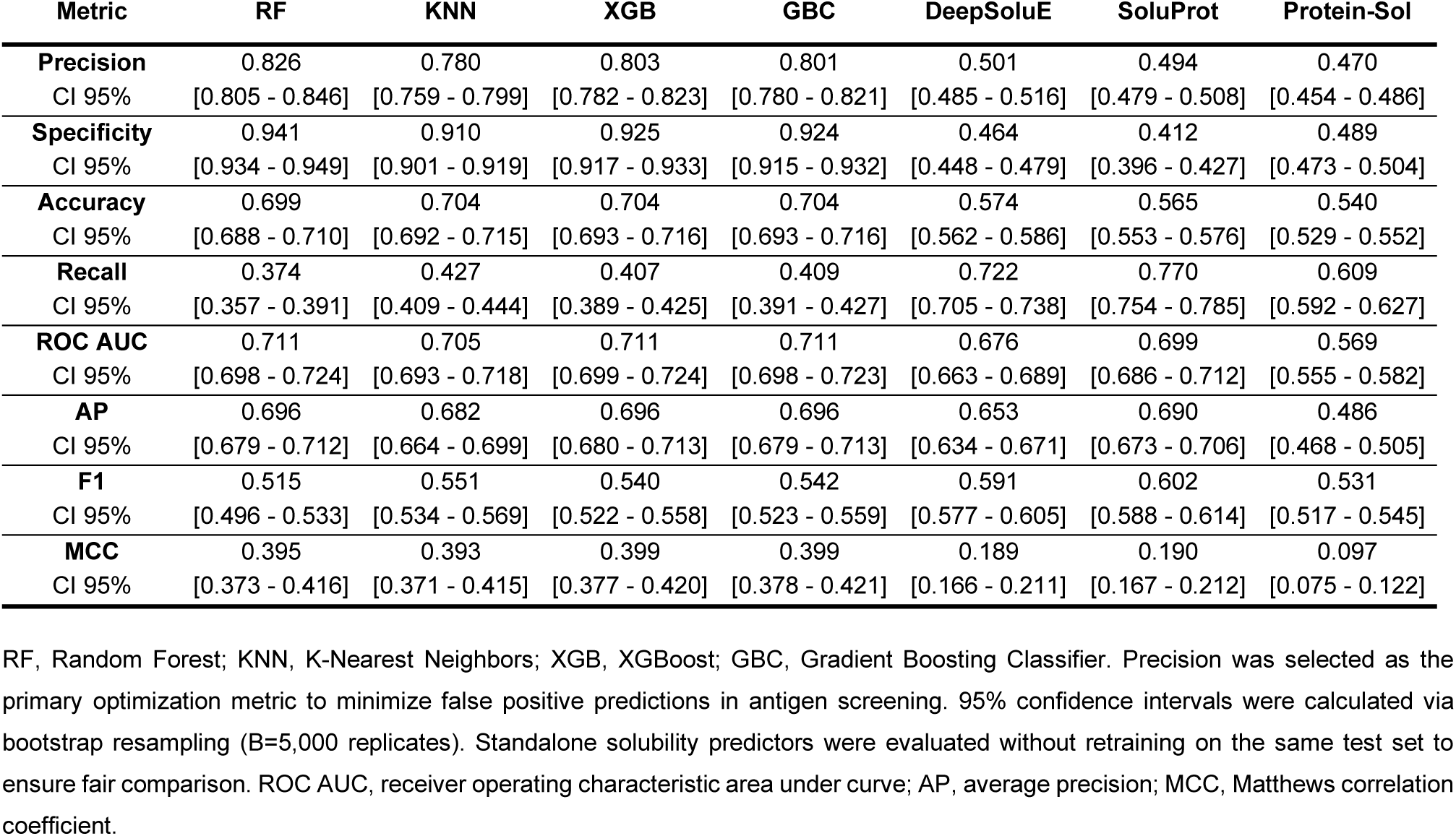
Performance comparison of supervised meta-learning models and individual solubility predictors on the held-out test set.

We next compared the best-performing ensemble model (RF) with the individual solubility predictors (Table 1, Fig 4D-F). The standalone predictors were not retrained and were evaluated directly on the same independent test set used for the supervised ensemble models, to ensure a consistent and fair comparison. RF achieved substantially higher precision (0.826, 95% CI [0.805–0.846]) compared to all three standalone tools: SoluProt (0.494), DeepSoluE (0.501), and Protein–Sol (0.470). For ROC-AUC and Average Precision, RF (0.711 and 0.696, respectively) showed performance comparable to or exceeding all standalone tools. Among the standalone predictors, SoluProt achieved comparable AP to RF (0.690 vs. 0.696, 95% CI [0.679–0.712]), but with substantially lower precision (0.494 vs. RF 0.826). RF’s superior precision and specificity (0.941 vs. 0.412), combined with equivalent ROC-AUC and AP, indicate that the ensemble provides more reliable antigen screening by minimizing false positive predictions.

To evaluate sequence identity-dependent effects, the RF test set was stratified into three identity ranges relative to the training set (Fig 5). The full test set (n=6,852) was divided into: samples with <30% identity (n=2,066), 30–80% identity (n=1,084), and >80% identity (n=447). Precision remained high and comparable across low and intermediate identity ranges (Fig 5A), with substantial decline at high sequence identity. ROC-AUC showed a similar pattern of performance degradation at high identity (Fig 5B), while AP maintained relatively high values across low and intermediate identity ranges before declining sharply at high identity (Fig 5C). These results indicate that RF achieved robust performance on sequences with low to moderate identity to the training set, while performance degraded substantially on sequences with high identity to training data, reflecting the presence of conflicting (high-identity, different-class) sequences within the training set.

**Fig 5.**
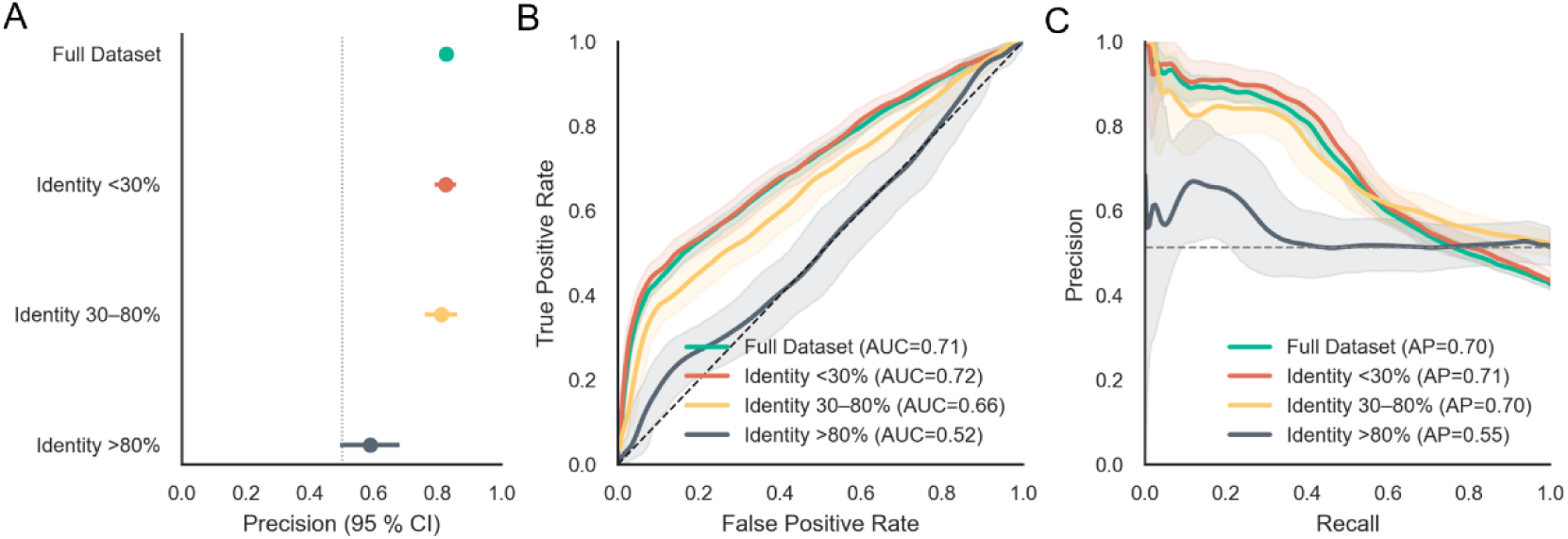
Performance of the Random Forest (RF) solubility meta-predictor on test sequences stratified by training set identity. The RF model was evaluated on test set samples stratified by sequence identity relative to the training set across four datasets: full test set (n=6,852), <30% identity (n=2,066, low identity to training), 30–80% identity (n=1,084, moderate-identity sequences), and >80% identity (n=447, high-identity sequences closely related to training data). **(A)** Precision with 95% bootstrap confidence intervals for each identity group. The vertical dashed line indicates the precision value for the full test set, serving as reference for comparison across identity-stratified subsets. **(B)** ROC curves with AUC values for each identity group. The black dashed diagonal line indicates random classifier performance (AUC=0.5). **(C)** Precision-Recall curves with Average Precision (AP) values for each identity group. The black dashed horizontal line indicates the baseline precision corresponding to the prevalence of soluble protein class in the full test set.

Permutation feature importance analysis was used to quantify the contribution of each standalone predictor to RF ensemble performance (Fig 6A-B). On the full test set, SoluProt more substantially contributed to both precision and ROC-AUC than ProtSol and DeepSoluE. The relative importance pattern remained consistent across identity ranges. SoluProt showed the largest contribution in the low-identity subset (<30%), followed by the intermediate-identity range (30–80%), with contributions substantially reduced in the high-identity subset (>80%). ProtSol and DeepSoluE contributed minimally to ensemble performance across all identity ranges. These results indicate that the ensemble relied predominantly on SoluProt while incorporating smaller, complementary contributions from the other predictors.

**Fig 6.**
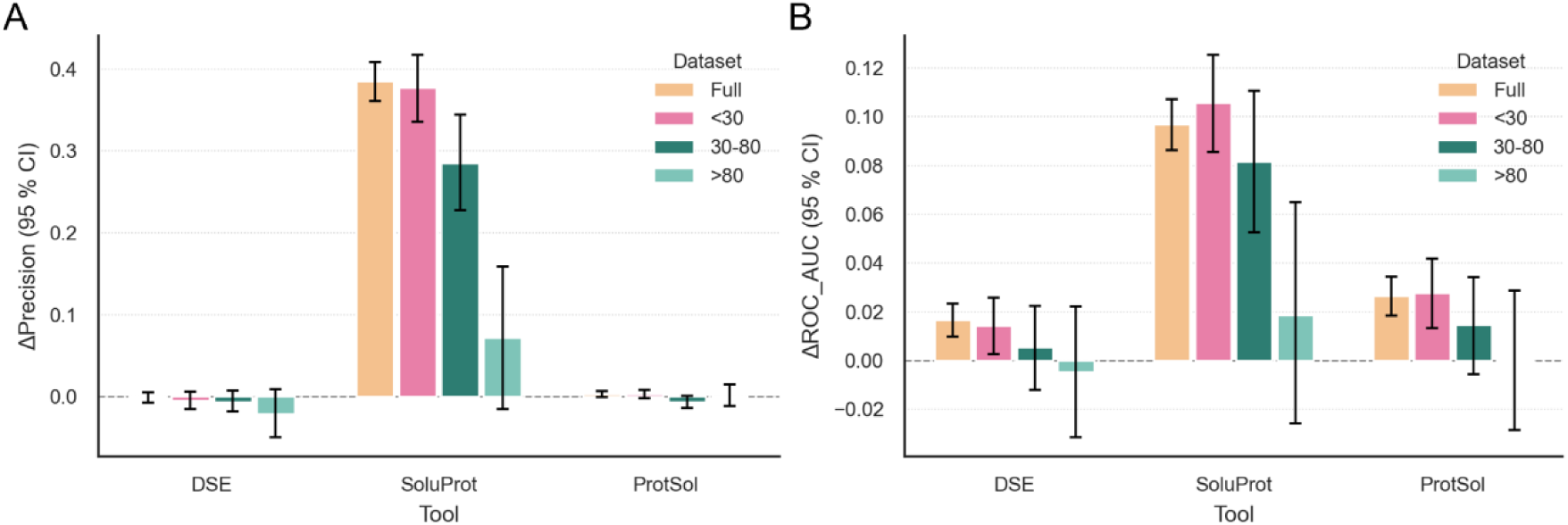
Permutation feature importance of solubility predictors in the RF meta-learner. Permutation feature importance analysis quantifies the contribution of each standalone solubility predictor (DeepSoluE, SoluProt, Protein–Sol) to the RF ensemble’s decision-making across four datasets: full test set, <30% identity to training, 30–80% identity to training, and >80% identity to training. (A) Change in precision (ΔPrecision) with 95% bootstrap confidence intervals for each predictor across datasets. (B) Change in ROC-AUC (ΔROC-AUC) with 95% bootstrap confidence intervals for each predictor across datasets.

#### Integrating stability predictions using supervised machine learning

Protein sequences with experimentally annotated thermal stability were collected from publicly available datasets associated with existing stability prediction tools. To minimize data leakage,, all proteins previously used to train the original predictors were removed.

Sequence redundancy was evaluated through pairwise identity analysis (Fig 7A). Most sequences clustered within the intermediate identity range (30–50%), accounting for approximately 75% of the dataset, whereas ∼18% showed low sequence identity (<30%). Only a minor fraction exhibited high sequence identity, indicating limited redundancy at stringent thresholds. Given the moderate redundancy and substantial sequence diversity, no redundancy reduction was applied. The final dataset comprised 20,777 proteins, including 8,904 thermophilic and 11,873 non-thermophilic sequences (Fig 7B, S4 Table).

**Fig 7.**
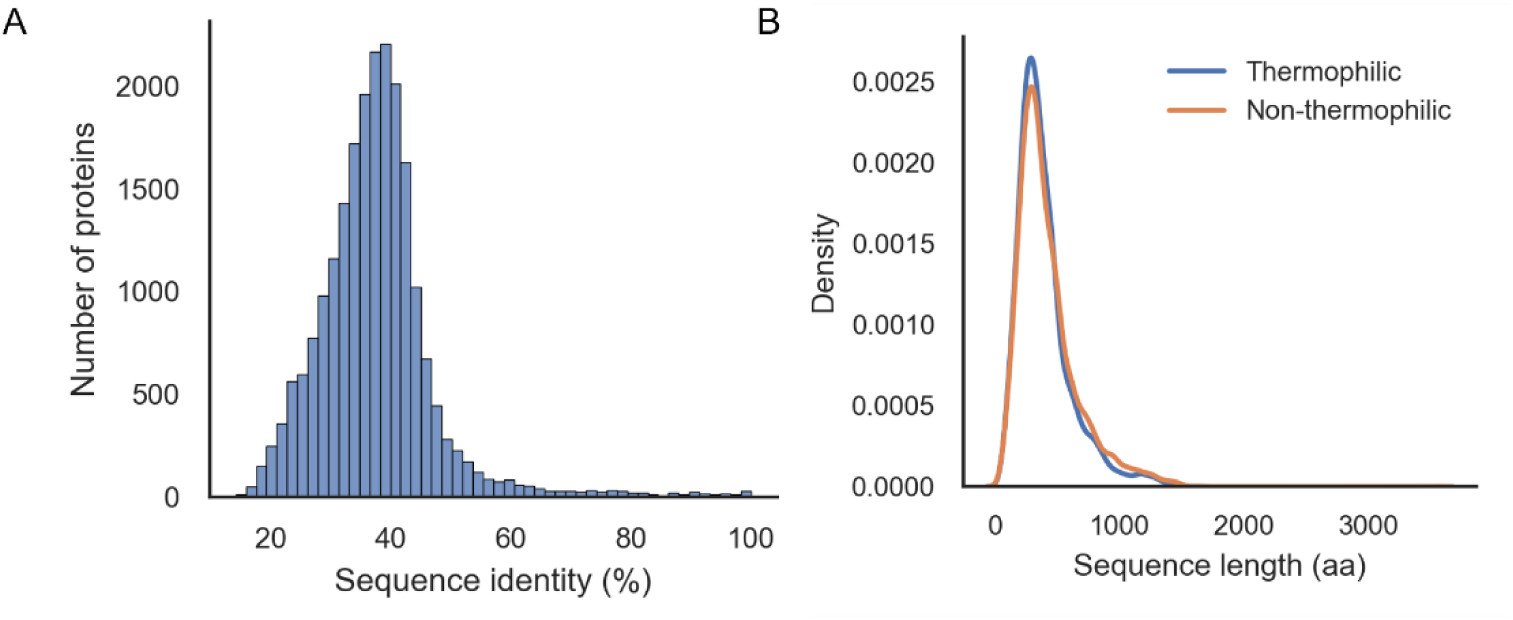
Sequence redundancy and length distribution in the thermal stability dataset. **(A)** Distribution of pairwise sequence identity. Most sequences cluster in the intermediate identity range (30–50%), with limited high-identity sequences (>70%). **(B)** Distribution of protein sequence lengths for thermophilic and non-thermophilic classes. The two classes show comparable length distributions.

Thermal stability scores generated by TemStaPro [23], ProLaTherm [24], and BertThermo [25] were integrated using four supervised meta-learning classifiers (RF, GBC, XGB, and KNN). Performance was evaluated on a held-out test set using precision as the primary optimization metric.

All four classifiers showed similarly high performance across evaluation metrics (Fig 8A-C). RF achieved the highest precision (0.961, 95% CI [0.951–0.970]), while all models displayed near-equivalent ROC-AUC (∼0.99) and AP values (∼0.98–0.99). Confidence intervals were estimated through bootstrap resampling (5,000 replicates). Based on its superior precision while maintaining equivalent discriminatory performance, RF was selected as the thermal stability meta-predictor.

**Fig 8.**
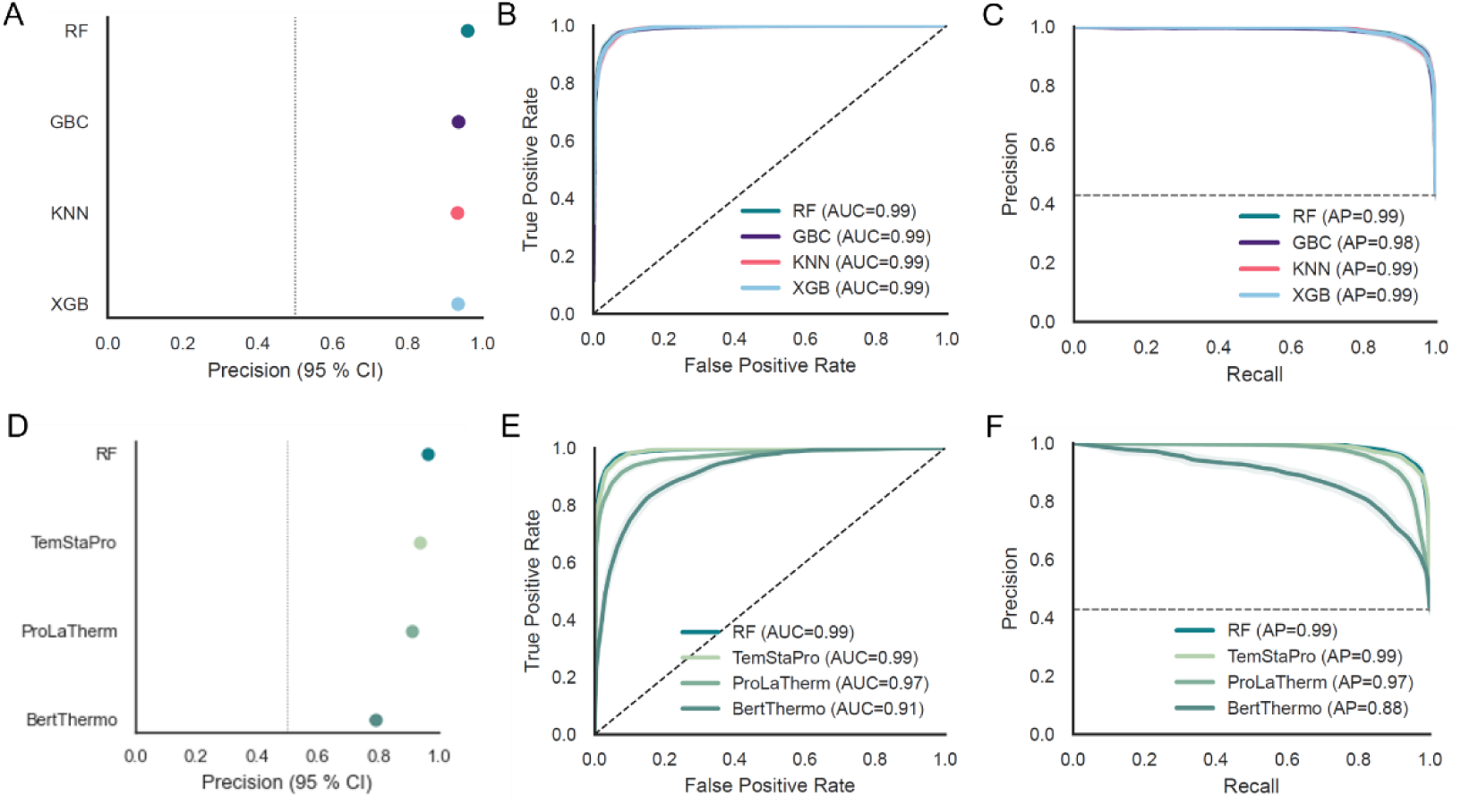
**Performance of supervised meta-learning models and comparison with individual thermal stability predictors**. **(A)** Precision values with 95% bootstrap confidence intervals for Random Forest (RF), Gradient Boosting Classifier (GBC), XGBoost (XGB), and K-Nearest Neighbors (KNN). **(B)** Receiver Operating Characteristic (ROC) curves showing threshold-independent discriminatory performance, with Area Under the Curve (AUC) values indicated. **(C)** Precision-Recall curves showing threshold-dependent ranking performance, with Average Precision (AP) values indicated. **(D)** Precision values with 95% bootstrap confidence intervals for RF and three standalone solubility predictors: Predictors: TemStaPro, ProLaTherm, and BertThermo. **(E)** ROC curves with AUC values for each method. **(F)** Precision-Recall curves with AP values for each method. Dashed lines indicate reference benchmarks: vertical (A, D: full test set precision), diagonal (B, E: random classifier, AUC=0.5), and horizontal (C, F: baseline precision).

We next compared the RF ensemble with the standalone thermal stability predictors on the same independent test set (Fig 8D-F, Table 2). RF achieved higher precision (0.961, 95% CI [0.951–0.970]) compared to all three standalone tools (Fig 8D). For ROC-AUC, RF showed performance comparable to TemStaPro while exceeding ProLaTherm and BertThermo (Fig 8E). Among the standalone tools, TemStaPro achieved the highest precision and ROC-AUC, approaching RF in both metrics. Precision-Recall curves showed similar ranking patterns across methods (Fig 8F).

**Table 2.**
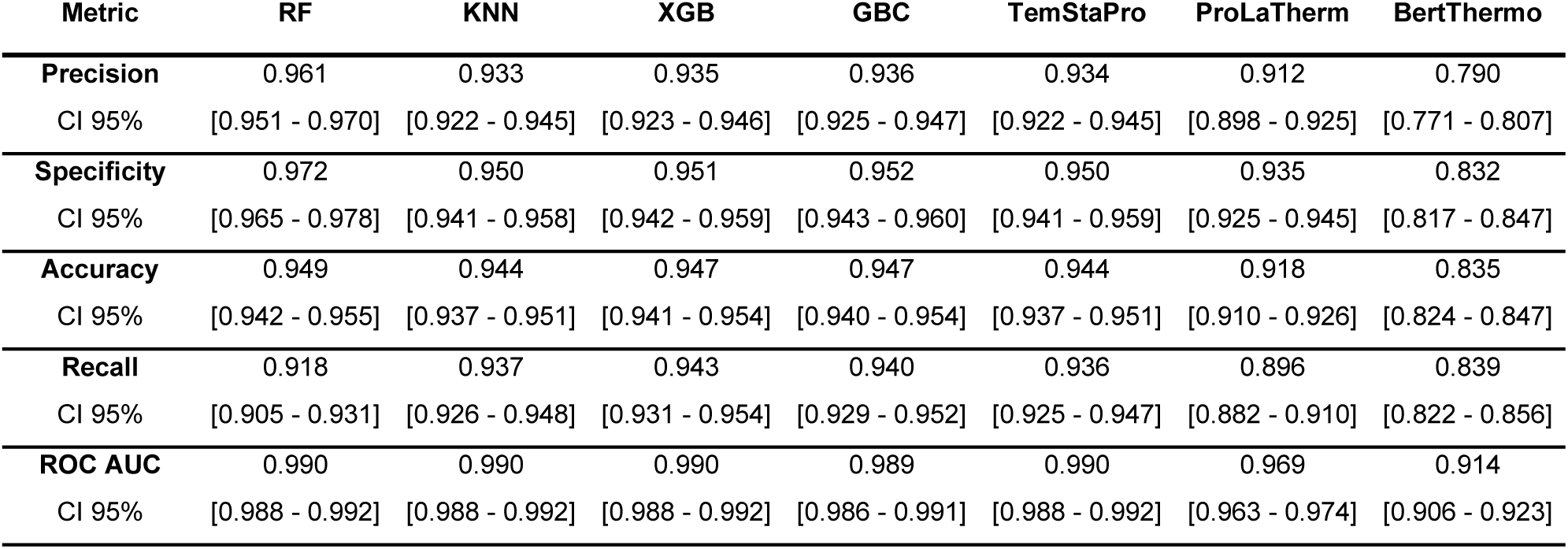
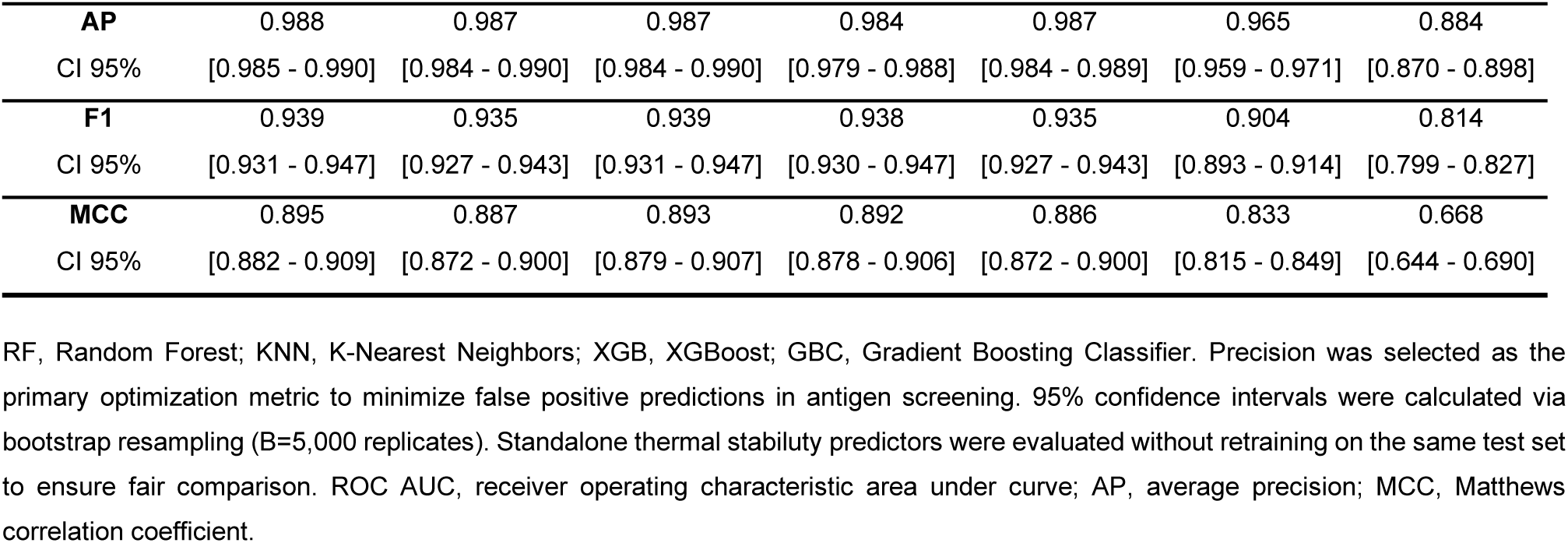
Performance comparison of supervised meta-learning models and individual thermal stability predictors on the held-out test set.

To evaluate whether RF thermal stability predictions showed sequence identity-dependent effects, the test set was stratified into three groups relative to training sequence identity (Fig 9A-C). The full test set comprised: sequences with <30% identity (n=462, minimal training overlap), 30–80% identity (n=3,350, substantial training overlap), and >80% identity (n=23, near-identical to training). Performance on low-identity sequences (<30%) remained high across all metrics. Precision achieved nearly 0.95 with comparable ROC-AUC (∼0.99) and AP (∼0.98), indicating robust generalization to divergent sequences. In the intermediate-identity range (30–80%), RF maintained consistently high performance with precision approaching 0.96 and ROC-AUC and AP both near 0.99, demonstrating strong predictive capability on this identity range. High-identity sequences (>80%) showed saturated performance with precision and AUC metrics reaching 1.0, indicating perfect predictions for sequences nearly identical to the training set.

**Fig 9.**
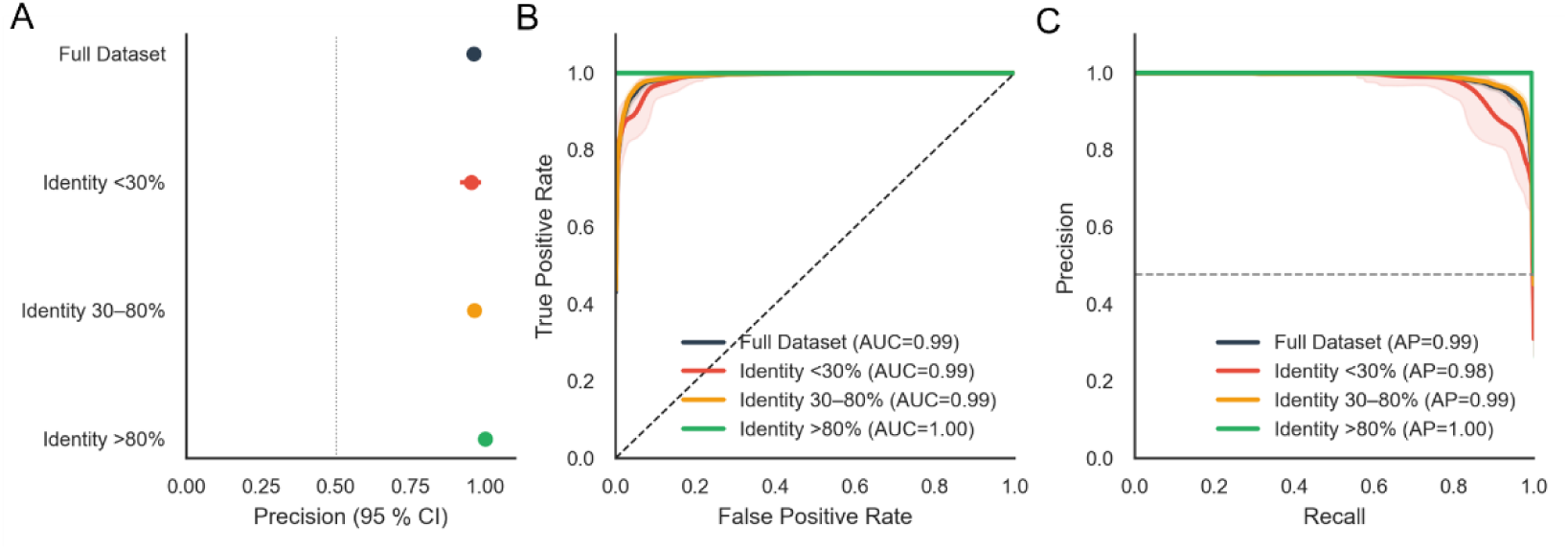
Performance of the Random Forest (RF) thermal stability meta-predictor on test sequences stratified by training set identity. The RF model was evaluated on test set samples stratified by sequence identity relative to the training set across four datasets: full test set, <30% identity (n=462, low identity to training), 30–80% identity (n=3,350, moderate identity), and >80% identity (n=23, high identity). **(A)** Precision with 95% bootstrap confidence intervals for each identity group. The vertical dashed line indicates the precision value for the full test set, serving as reference for comparison across identity-stratified subsets. **(B)** ROC curves with AUC values for each identity group. The black dashed diagonal line indicates random classifier performance (AUC=0.5). **(C)** Precision-Recall curves with Average Precision (AP) values for each identity group. The black dashed horizontal line indicates the baseline precision corresponding to the prevalence of thermophilic class in the full test set.

Permutation feature importance analysis revealed differential contributions of the three thermal stability predictors to the RF ensemble (Fig 10A-B). TemStaPro provided substantially larger marginal signal to both precision and ROC-AUC across all identity ranges, while ProLaTherm and BertThermo contributed at smaller scales. The RF ensemble achieved precision improvements over TemStaPro alone (0.9606 vs 0.9339, approximately 2.7 percentage points), despite the unbalanced importance contributions. This pattern suggests that while TemStaPro captured the primary thermal stability signal, ProLaTherm and BertThermo provided complementary information that, although modest, contributed to incremental performance gains. The consistency of this pattern across sequence identity ranges, indicating robust performance independent of training set composition.

**Fig 10.**
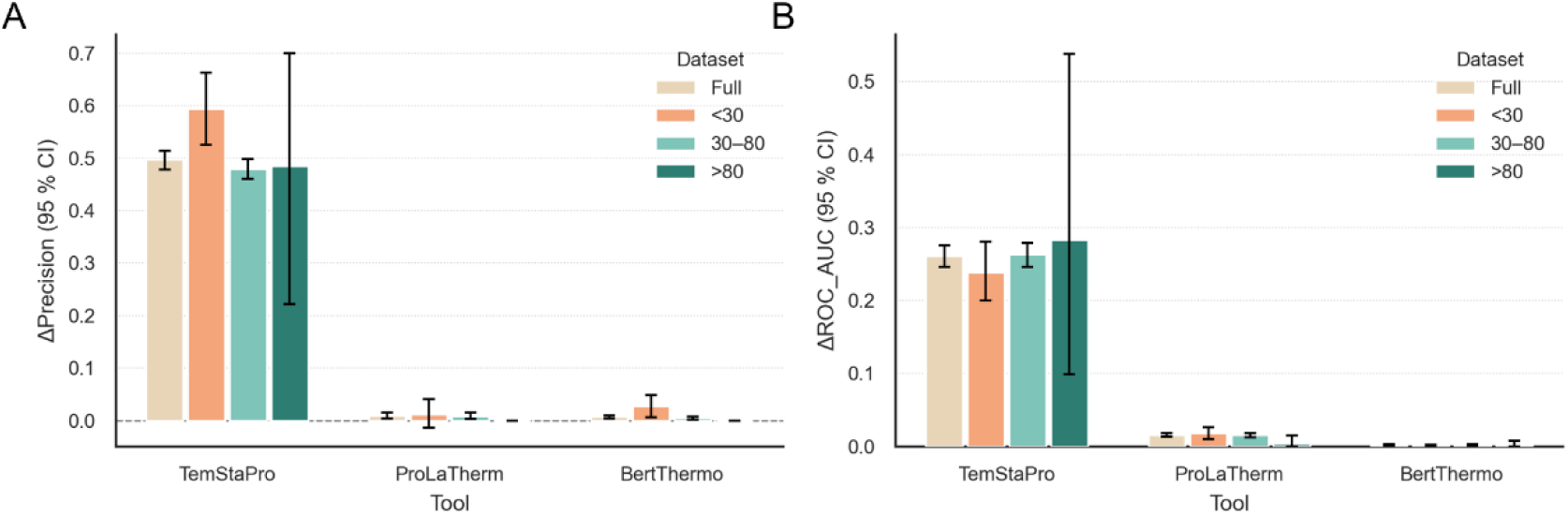
Permutation feature importance of thermal stability predictors across sequence identity ranges. Permutation feature importance analysis quantifies the contribution of each standalone thermal stability predictor (TemStaPro, ProLaTherm, and BertThermo) to the RF ensemble across four datasets: full test set, <30% identity to training, 30–80% identity to training, and >80% identity to training. **(A)** Change in precision (ΔPrecision) with 95% bootstrap confidence intervals for each predictor. **(B)** Change in ROC-AUC (ΔROC-AUC) with 95% bootstrap confidence intervals for each predictor.

#### Integration of solubility and stability into the Expression Efficiency Score

To examine the integrated developability metric, the EES was applied to an in-house panel of 36 recombinant proteins expressed in *E. coli* from diverse biological sources (28 soluble, 8 insoluble, S5 Table). Notably, solubility classification in this validation set was defined by subcellular localization (soluble cytoplasm vs. inclusion bodies) rather than the ultracentrifugation-based annotations used in model training, enabling assessment across distinct experimental definitions of solubility.

Soluble proteins exhibited higher mean predicted solubility scores than insoluble proteins (0.397 ± 0.243 vs. 0.270 ± 0.224; Cliff’s δ = 0.348, medium effect size). Predicted stability scores were also moderately higher in soluble proteins (0.148 ± 0.254 vs. 0.059 ± 0.066), although the associated effect size was negligible (Cliff’s δ = −0.098),. Among the evaluated metrics, EES showed the strongest class separation, with mean values of 0.347 ± 0.187 for soluble proteins and 0.228 ± 0.174 for insoluble proteins (Cliff’s δ = 0.366, medium effect size).

Although statistical significance was not achieved (likely due to the limited number of insoluble proteins, n = 8), the observed effect sizes support the biological relevance of the integrated score. Because the validation dataset lacked direct experimental measurements of thermal stability, solubility represented the only experimentally validated phenotype. Consistently, proteins with higher EES values clustered predominantly within the high-solubility region of the solubility–stability space (Fig 11), supporting the stronger weighting assigned to solubility (0.8) in the EES formulation. These results validate the EES as a practical ranking tool for prioritizing protein constructs based on expression efficiency predictions, with solubility predictions serving as the primary determinant of heterologous expression success in mesophilic *E. coli* production systems.

**Fig 11.**
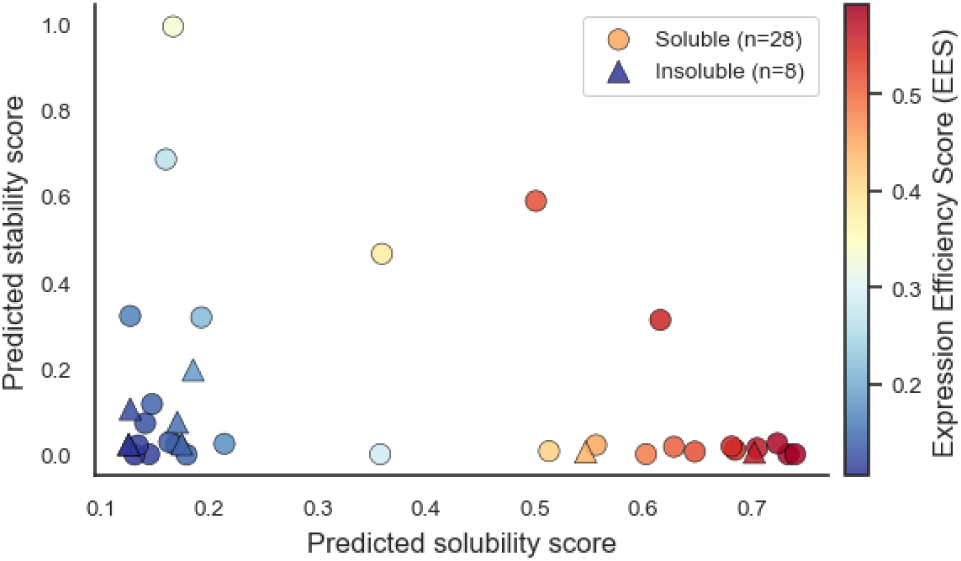
Validation of Expression Efficiency Score (EES) predictions for solubility and thermal stability. Independent validation of 36 recombinant proteins from diverse biological sources. The ESS is depicted by color intensity, ranging from blue (low expressivity) to red (high expressivity). Soluble proteins are shown as circles (n=28) and insoluble proteins as triangles (n=8).

## Discussion

Advances in sequencing and structural modeling have shifted antigen discovery from data scarcity to decision complexity. Since candidate lists have greatly expanded, the challenge has moved toward prioritizing antigenic proteins using reproducible and biologically grounded criteria. BIOINF-farma was developed to address this need by integrating epitope prediction with developability assessment in heterologous expression systems, enabling comparative evaluation of candidates within a single computational framework.

A distinctive feature of BIOINF-farma is the integration of the entire workflow, from raw sequence processing and structure acquisition/prediction to antigenicity and developability scoring, within a single computational pipeline operating in a consistent environment. Importantly, the pipeline accepts either protein sequences or three-dimensional structures as input, making the platform independent of the initial data format. This flexibility removes upstream constraints on the format of available data and simplifies adoption across heterogeneous datasets. This capability is enabled by the initial module, which also supports de novo three-dimensional structure prediction. This design removes the manual handoffs and format conversions that typically fragment analyses across separate tools and makes whole-proteome screening operationally feasible in the context of emerging pathogen responses.

A defining limitation of current B-cell epitope prediction workflows is the coexistence of two fundamentally different predictor types: sequence-based tools, such as BepiPred 3.0, provide residue-level linear epitope propensity scores [19,28], whereas structure-based methods identify conformational surface epitopes [11,17,29,30]. These approaches are highly informative for mapping potential antibody-binding regions, but they do not allow to produce a standardized score that allows direct comparison between whole proteins. Some existing frameworks, notably VaxiJen, assign protein-level antigenicity scores through alignment-independent physicochemical descriptors [31]. However, such approaches typically operate on primary sequence and do not explicitly incorporate three-dimensional structural information, thereby missing conformational epitopes, whose spatial accessibility depend on native protein folding. In BIOINF-farma, we addressed this limitation by systematically integrating structure-based (MLCE/REBELOT-BEPPE) and sequence-based (BepiPred 3.0) epitope predictions through residue-level combination and protein-level aggregation. This approach allowed us to construct an Antigenicity Score that captures both linear and conformational epitope signals. Quantitative validation on the manually curated Antigenicity Dataset (CAD, n=190 proteins) demonstrated marked score separation between likely antigenic and non-antigenic classes, validating the utility of the integrated signal for binary classification.

While antigenicity is central to vaccine candidate selection, it is equally important that antigens can be efficiently expressed in heterologous systems, yet these two aspects are rarely evaluated together. In most computational workflows, immunogenic potential and recombinant feasibility are treated as separate problems. In contrast, BIOINF-farma explicitly integrates these dimensions through a developability module, summarized in the EES, which combines solubility and thermal stability predictions.

Protein solubility remains one of the most challenging properties to predict from sequence. Soluble expression in heterologous systems such as *E. coli* depends not only on intrinsic amino acid composition, but also on folding kinetics, aggregation propensity, translation rate, chaperone availability, and experimental conditions. As highlighted in methodological reviews, current solubility predictors typically achieve only moderate performance, reflecting the multifactorial and context-dependent nature of the phenotype [12,20–22,32]. Supervised meta-learning through Random Forest integration substantially improved precision, with a 65% relative improvement over the best individual tool. Critically, this improvement was accompanied by maintained or superior ranking performance (ROC-AUC and AP), indicating that the ensemble refined classification boundaries without compromising overall discriminatory capacity. Precision gain reflects a substantial reduction in false positive predictions, enhancing the reliability of antigen screening by ensuring that candidates predicted as soluble are more likely to express successfully and concentrating experimental resources on higher-confidence leads. The performance gain arose from complementary information integration: permutation analysis revealed that SoluProt contributed the largest marginal signal to both precision and ROC-AUC, while DeepSoluE and Protein–Sol provided smaller, non-redundant contributions. Interestingly, RF performance degraded substantially at high sequence identity to the training set (>80%), reflecting the presence of conflicting high-identity examples within the training data rather than inherent ensemble limitations. This identity-dependent behavior highlights an important consideration for model deployment: generalization on sequences with low-to-moderate identity to training data (<80%) was robust, while performance on high-identity sequences became unreliable, likely reflecting unresolved biological variability within these similar clusters.

Thermal stability prediction presents a different scenario. The relationship between sequence features and thermophilicity is comparatively stronger and more systematic, reflecting well-characterized structural adaptations in thermostable proteins, such as enhanced ionic networks and optimized packing interactions [9]. Accordingly, individual stability predictors already demonstrated strong baseline performance [23–25]. Nevertheless, supervised ensemble integration provided additional gains: RF achieved higher precision, corresponding to a ∼2.7 percentage point improvement over the best individual tool, while maintaining equivalent ROC-AUC (∼0.99). Notably, this performance remained consistent across sequence identity ranges, indicating stable ensemble behavior regardless of training set identity, a feature particularly relevant for predicting thermophilicity of novel organisms or divergent sequence contexts.

The divergent performance profiles of solubility and stability meta-learners highlight an important principle in ensemble design: integration strategies should be calibrated to the underlying signal strength and heterogeneity of the problem domain. In solubility prediction, where uncertainty is substantial and individual tools exhibit systematic biases, ensemble integration provides significant bias mitigation and precision improvements. In thermal stability, where individual predictors are already strong, ensemble integration functions primarily as a calibration mechanism, refining decision thresholds while contributing modest incremental gains. In both cases, extensive refinement of training data and evaluation on independent, identity-stratified test sets reduced risk of circularity and provided more realistic generalization estimates than benchmark evaluations typically report.

To evaluate the integrated developability metric, EES predictions were validated on an independent panel of 36 recombinant proteins expressed in *E. coli*. Soluble proteins exhibited significantly higher average EES than insoluble counterparts, demonstrating meaningful biological discrimination despite the limited insoluble sample size (n=8). Notably, solubility and thermal stability contributed independently to this differentiation: soluble proteins showed elevated mean solubility and stability scores. The prevalence of high-EES scores (>0.45) in the high-solubility region validated the primary weighting of solubility (0.8) in the EES formulation. Exceptions like the archaeal protein StLASPO [33], which achieved exceptional predicted stability (0.995) but low solubility (resulting in moderate EES=0.332), illustrate how independent assessment of these properties captures complementary aspects of expression phenotypes and prevents over-reliance on single determinants.

Several limitations warrant consideration. First, de novo structure prediction via Boltz-2 yields monomeric models only; consequently, antigenicity predictions cannot account for quaternary organization and its effects on epitope accessibility in multimeric native proteins. Second, the integration of glycosylation patterns is not currently implemented; glycan shielding can modulate epitope accessibility in ways not captured by structure-based or sequence-based prediction alone[34]. Third, although supervised integration improves robustness, predictive performance ultimately depends on the representativeness of available experimental datasets, which remain biased toward naturally occurring proteins. Finally, our validation against subcellular localization (soluble cytoplasm vs. inclusion bodies) differs from the ultracentrifugation-based classification used in training. Future work should evaluate transfer across distinct solubility phenotypes more systematically and could include training on context-specific datasets (e.g., organism-, protein family-, or application-specific), to enable more tailored and accurate predictions for defined use cases.

Conceptually, BIOINF-farma illustrates a broader principle that is becoming increasingly relevant in computational biology as biological datasets continue to expand: the transition from isolated prediction modules toward integrated, interpretable frameworks that jointly optimize multiple objectives. Antigen selection inherently requires balancing immunogenic potential against expression feasibility, a trade-off that cannot be adequately addressed through independent optimization of separate objectives. By providing clear, benchmarkable protein-level scores integrating diverse computational signals, BIOINF-farma offers a more reproducible and operationally integrated framework for antigen discovery.

## Availability and future directions

BIOINF-farma is freely accessible through a web-based graphical interface at www.bioinf-farma.uninsubria.it/ and as an open-source Python/bash pipeline available on GitHub (https://github.com/HhrBnd/bioinf-farma.git, DOI: 10.5281/zenodo.20156208) under the GNU Affero General Public License (AGPL-3.0-or-later). The complete source code, including pre-trained Random Forest meta models, conda environment specifications, and curated datasets, is provided through the repository and/or supplementary materials. The pipeline was developed using modular scripts and a user-friendly web interface to ensure accessibility for experimental immunologists without requiring programming expertise. The platform graphical user interface enables remote access for rapid evaluation of protein candidates without requiring programming expertise. The graphical user interface enables remote evaluation of protein candidates without the need for local computational resources, facilitating integrated antigen prioritization for researchers without specialized bioinformatics training. This accessibility may be particularly valuable in vaccine research during pathogen emergencies, where rapid decision-making and candidate prioritization are critical.

Future developments will focus on expanding the scope, accuracy, and applicability of BIOINF-farma. Key priorities include: (i) enlarging the meta-learner training datasets to improve coverage of underrepresented organisms and protein families; (ii) extending developability assessment to alternative heterologous systems, including plant-based, mammalian, and insect cell platforms); (iii) integrating glycosylation prediction and post-translational modification modeling; (iv) incorporating T-cell epitope prediction to complement the current B-cell-oriented framework; and (v) adding phylogenetic and genomic context analysis to assess cross-reactivity risks and host immune escape potential. Community feedback and external contributions are encouraged through the GitHub repository, fostering continuous improvement and long-term sustainability of the platform.

## Materials and Methods

### Curated antigenicity dataset

A manually curated reference dataset, the Curated Antigenicity Dataset (CAD), was assembled to evaluate antigenicity predictions. Immunogenic protein sequences were collected from publicly available immunogenicity prediction resources [31,35,36], and corresponding three-dimensional structures were retrieved from the RCSB Protein Data Bank when available [37]. Additional structures of well-characterized viral and bacterial antigens were obtained from the Immune Epitope Database [4], whereas, PDB structures of human non-immunogenic proteins were obtained from the Tumor Immunogens Database (www.ddg-pharmfac.net/vaxijen3/tumordb/home/) and integrated with structures of abundant human serum proteins. Only PDB structures with resolution ≤ 3.5 Å and without missing residues in structured regions were selected to ensure compatibility with Amber-based preprocessing and structure-based epitope prediction using MLCE/REBELOT-BEPPE. PDB files were manually curated based on biological assembly annotations to preserve biologically relevant structural organization, retaining all chains for oligomeric proteins and splitting entries into independent functional entities when appropriate. Functional information, cellular localization, and evidence of immune exposure were evaluated by consulting UniProt (Feature Viewer)[38]. Based on curated biological criteria, proteins were classified as likely antigenic or likely non-antigenic for validation purposes.

### Antigenicity score threshold optimization

To define an optimal decision threshold for the binary classification of proteins as antigenic or non-antigenic, sensitivity and specificity were evaluated across score thresholds on the CAD using the Youden’s J statistic (J = sensitivity + specificity − 1), which maximizes balanced accuracy [39]. The resulting threshold represents a dataset-calibrated operating point. Statistical differences in antigenicity score distributions between antigenic and non-antigenic proteins were assessed using the Mann–Whitney U test, a non-parametric method that does not assume normality of the underlying distributions [40]. The magnitude of the observed effect was quantified using Cliff’s delta, a non-parametric effect size measure estimating the probability that antigenicity scores from antigenic proteins exceed those from non-antigenic proteins [41]. All statistical analyses were performed in Python using SciPy and scikit-learn libraries[42].

### Solubility reference dataset construction and curation

Multiple sequence-based protein solubility datasets were integrated to construct the solubility dataset used in this study. These included datasets released with DeepSoluE [20], SoluProt [21], and Protein–Sol [22], as well as publicly available solubility datasets derived from experimental studies of recombinant protein expression, including SoDoPe [43] eSOL [44] and NESG datasets [45]. To prevent data leakage and circularity, all protein sequences used for training the individual solubility predictors were first merged into a unified reference set and subsequently removed from the complete collection of available solubility data. Only protein sequences with unambiguous experimental solubility annotations were retained.

### Stability reference dataset construction and curation

Protein sequences annotated with experimental information on thermal stability were collected from publicly available *E. Coli*-related datasets associated with existing stability prediction tools, including BertThermo [25], DeepTP [46], iThermo [47], ProLaTherm [24], and SAPPHIRE [48]. Due to substantial overlap among these resources, all protein sequences used for training BertThermo [25], ProLaTherm [24], and TemStaPro [23] were aggregated into a unified reference set and removed from the complete collection to generate an unbiased dataset. Only protein sequences with unambiguous experimental annotations were retained. Thermal stability was treated as a binary classification task according to the labeling schemes reported in the original datasets.

### Sequence redundancy assessment

MMseqs2 was used to perform all-versus-all sequence comparisons independently across both the solubility and stability datasets [14]. For each protein, the highest sequence identity to any other sequence in the dataset (excluding self-matches) was retained to characterize redundancy. Based on these analyses, different strategies were adopted for the two datasets. For the stability dataset, no redundancy reduction was applied, and the full dataset was retained for downstream analyses. For the solubility dataset, redundancy reduction was performed separately within each class (soluble and insoluble). Protein sequences were clustered using MMseqs2 at a sequence identity threshold of 25%, and a single representative sequence was retained from each cluster. This procedure was applied independently to the two classes, generating non-redundant subsets of soluble and insoluble proteins. The resulting datasets were used for subsequent model development and evaluation.

### Meta-learner development and training

Solubility scores were generated using DeepSoluE, SoluProt, and Protein–Sol [20–22]. Thermal stability scores were obtained using BertThermo, ProLaTherm, and TemStaPro [23–25]. These tools were selected based on public availability, redistributable licensing, transparency of training data, and compatibility with local installation within a controlled computational environment. The objective of this module is to derive robust protein-level developability estimates through supervised meta-learning applied to multiple predictor outputs.

All predictors were executed using their official releases and default configurations. Outputs from individual predictors were used as input features for supervised integration. Models were implemented in Python (v3.12.2) using scikit-learn (v1.4.1) [42]. Datasets were partitioned into training and test sets using an 80/20 stratified split. Hyperparameter tuning was conducted via grid search with repeated stratified 5-fold cross-validation (5 folds × 3 repeats; 15 validation splits). Four classifiers were evaluated: k-nearest neighbors (KNN), Random Forest (RF), Gradient Boosting (GB), and Extreme Gradient Boosting (XGBoost) [49]. Precision was selected as the primary optimization metric to minimize false positive predictions in antigen screening workflows.

### Performance evaluation and model interpretation

Model performance was assessed on the held-out test set using ranking-based and decision-level metrics. Primary evaluation focused on ranking performance, assessed using the receiver operating characteristic (ROC) curve and the precision–recall curve. ROC-AUC (area under the ROC curve) provides a threshold-independent measure of the model’s ability to discriminate between classes across all decision thresholds. Average Precision (AP) was computed from the precision–recall curve to assess ranking performance. For completeness, additional decision-level metrics were computed at a fixed threshold of 0.5, including precision, recall (sensitivity), specificity, accuracy, F1 score, and Matthews correlation coefficient (MCC).

To quantify statistical uncertainty, 95% confidence intervals (CI) were estimated using non-parametric bootstrap resampling of the held-out test set (B=5,000 replicates). In each replicate, test samples were drawn with replacement and performance metrics recomputed. The 2.5th and 97.5th percentiles of the resulting bootstrap distributions were used to define confidence intervals. Bootstrap resampling was applied only to the test set and was not used during cross-validation.

### Identity-stratified generalization analysis

To assess generalization as a function of sequence identity to the training set, test sequences were stratified by sequence identity relative to the training set using MMseqs2 [14]. For each test sequence, the best-hit fractional identity (fident) was used to assign the sequence to one of three identity ranges: <30% identity (low-identity sequences, divergent from training data), 30–80% identity (moderate-identity sequences, intermediate distance from training sequences), and >80% identity (high-identity sequences, closely related to training data). Performance metrics were computed separately within each identity range. This analysis provides an estimate of predictive performance as a function of sequence identity to the training dataset and helps identify potential residual redundancy effects on model generalization [50].

### Permutation feature importance

To assess the contribution of individual predictors within the selected meta-model, permutation feature importance was computed on the held-out test set [51,52]. For each predictor, its values were randomly permuted across samples while keeping all other inputs unchanged. The resulting decrease in model performance (ΔPrecision and ΔROC-AUC) relative to the original model was measured. Larger performance drops indicate greater reliance on that predictor. Permutation importance quantifies how much each predictor contributes to model predictions. A large decrease in performance when permuting a predictor indicates that the model relies heavily on that input, either because it carries a unique signal or because it captures information redundant with other correlated predictors. Permutation importance was computed on the full test set and separately within the three test vs train data identity ranges previously described (i.e.: < 30%, 30-80%, >80%).

### In-house validation panel

To assess the integrated score in an applied setting, an independent in-house dataset comprising 36 recombinant proteins expressed in *E. coli* was analyzed. For each construct, the experimentally observed expression outcome was recorded based on the reported soluble fraction, distinguishing proteins that yielded predominantly soluble material, predominantly inclusion bodies, or condition-dependent solubility. For all proteins in this panel, solubility and stability scores were computed using the corresponding meta-models, and the EES was derived as described above. This dataset was not used for model training or hyperparameter selection.

## Acknowledgements

We thank Dr. Federica Taddio for her assistance in software testing and prof. Simone Tini for helpful insights into the design of the graphical interface.

## Supporting information

S1 Text. Installation instructions and a tutorial.

S2 Table. Curated Antigenicity Dataset (CAD). The dataset comprises 190 manually annotated proteins (115 likely antigenic, Label = 1; 75 likely non-antigenic, Label = 0) collected from IEDB, VaxiJen, Tumor Immunogens Database, and Immunohub. Columns include UniProt accession, protein name, organism, PDB identifier with chain designation, classification label, sequence length, and computed antigenicity score (S_protein, 0–1 range). All structures have resolution ≤ 3.5 Å and no missing residues.

S3 Table. Curated Solubility Dataset. The dataset comprises 34,476 non-redundant protein sequences (14,770 soluble, 19,706 insoluble) integrated from DeepSoluE, SoluProt, Protein-Sol, SoDoPe, eSOL, and NESG databases, with all training sequences removed to prevent data leakage. Columns include protein identifier, amino acid sequence, sequence length, source dataset, and experimental solubility label (Solubility_exp: 1 = soluble, 0 = insoluble). Used for training and validation of solubility meta-learner.

S4 Table. Thermal Stability Dataset. The dataset comprises 20,777 protein sequences (8,904 thermophilic, 11,873 non-thermophilic) integrated from BertThermo, DeepTP, iThermo, ProLaTherm, and SAPPHIRE databases, with all training sequences removed to prevent data leakage. Columns include amino acid sequence, protein identifier, and experimental binary classification (exp_binary: 1 = thermophilic, 0 = non-thermophilic). Used for training and validation of thermal stability meta-learner.

S5 Table. Independent In-house Validation Panel. The validation dataset comprises 36 recombinant proteins expressed in E. coli from diverse biological sources (28 soluble in cytoplasm, 8 insoluble in inclusion bodies) used for independent evaluation of BIOINF-farma predictions. Columns include protein identifier, predicted solubility and stability scores (0–1 range), corresponding binary predictions (0 = negative, 1 = positive), Expression Efficiency Score (S_EES), organism source, expression host, experimental validation status (experimental_clean: YES/NO indicating confirmed expression phenotype), and amino acid sequence. Solubility phenotype defined by subcellular localization rather than ultracentrifugation-based classification used in model training, enabling assessment of generalization across distinct solubility phenotypes.

